# Detecting effective starting point of genomic selection by divergent trends from BLUP and ssGBLUP in pigs, beef cattle, and broilers

**DOI:** 10.1101/2021.05.28.446145

**Authors:** Rostam Abdollahi-Arpanahi, Daniela Lourenco, Ignacy Misztal

## Abstract

Genomic selection has been adopted nationally and internationally in different livestock and plant species. However, understanding whether genomic selection has been effective or not is an essential question for both industry and academia. Once genomic evaluation started being used, estimation of breeding values with pedigree BLUP became biased because this method does not consider selection using genomic information. Hence, the effective start point of genomic selection can be detected in two possible ways including the divergence of genetic trends and Realized Mendelian sampling (RMS) trends obtained with BLUP and Single-step genomic BLUP (ssGBLUP). This study aimed to find the start date of genomic selection for a set of economically important traits in three livestock species by comparing trends obtained using BLUP and ssGBLUP. For this purpose, three datasets comprised a pig dataset with 117k genotypes and 1.3M animals in pedigree, Angus cattle dataset consisted of ~842k genotypes and 11.5M animals in pedigree, and a purebred broiler chicken dataset included ~154k genotypes and 1.3M birds in pedigree were used. The genetic trends for pigs diverged for the genotyped animals born in 2014 for average daily gain and backfat. In beef cattle, the trends started diverging in 2009 for weaning weight and in 2016 for postweaning gain, with little diverging for birth weight. In broiler chickens, the genetic trends estimated by ssGBLUP and BLUP diverged at breeding cycle 6 for two out of three production traits. The RMS trends for the genotyped pigs diverged for animals born in 2014, more for average daily gain than for backfat. In beef cattle, the RMS trends started diverging in 2009 for weaning weight and in 2016 for postweaning gain, with a trivial trend for birth weight. In broiler chickens, the RMS trends from ssGBLUP and BLUP diverged strongly for two production traits at breeding cycle 6, with a slight divergence for another trait. Divergence of the genetic trends from ssGBLUP and BLUP indicates onset of the genomic selection. Presence of trends for RMS indicates selective genotyping, with or without the genomic selection. The onset of genomic selection and genotyping strategies agree with industry practices across the 3 species. In summary, the effective start of genomic selection can be detected by the divergence between genetic and RMS trends from BLUP and ssGBLUP.

## Introduction

Genomic selection has been widely recognized as a successful tool for genetic improvement, as evident by the extensive genotyping in various livestock and plant species (Misztal et al., 2020; VanRaden, 2020). Genomic selection allows to preselect young animals and also parents with higher accuracy than with BLUP (Patry and Ducrocq, 2011a; Tyrisevä et al., 2018b). However, the actual gains with genomic selection depend on a number of factors, aside from the genetic parameters. These include the choice of animals for genotyping, quality of methods for genomic prediction, and fraction of genotyped animals used for breed improvement. Genotyping is not effective if only parents with large number of progenies are genotyped because their BLUP evaluations are already accurate. A genomic selection scheme using simple single-trait models, possibly with few phenotypes, may be less accurate than BLUP selection with more complete data and models (Muir, 2007). Finally, if genotyping is used only for marketing, e.g., young bull sales to commercial farms, such genotyping has no effect on the genetic improvement of the breeding population.

With a large investment in genomic selection, it is of interest to find out the onset of the genomic selection and whether it is successful over the long run. There are several possible ways to find out the start date of genomic selection. One way to investigate the onset of genomic selection is by analyzing differences in genetic trends by BLUP and single-step genomic BLUP (ssGBLUP). Under genomic selection, BLUP cannot account for the fact that animals are being selected based on genomic information before having their phenotypes recorded (i.e., genomic preselection) and is therefore biased (Party and Ducrocq, 2009; Patry and Ducrocq, 2011b). On the contrary, ssGBLUP accounts for all sources of information jointly and is expected to be less affected by preselection bias (Legarra et al., 2009; VanRaden and Wright, 2013; Legarra et al., 2014). Superior genetic trends by ssGBLUP compared to BLUP have been reported in several cases. Masuda et al. (2018) presented trends for milk yield in Holsteins by BLUP and ssGBLUP. While the trend by ssGBLUP increased at the expected beginning of the genomic selection, the trend by BLUP leveled off. Koivula et al. (2018) reported that including the genotypes of culled bull calves in the ssGBLUP analysis leads to higher genetic trends for milk production traits of Nordic Red Dairy Cattle compared to the situation where genomic information of the culled bull calves is ignored.

Another way to investigate the onset of genomic selection is by analyzing genetic and phenotypic trends, expecting accelerating trends under genomic selection (Misztal et al., 2020). However, both trends are affected by changes in selection policies and incur some lag time. Additionally, changes in genetic parameters over time (Hidalgo et al., 2020) may cause fluctuations in the genetic trend. The third way is by analyzing realized Mendelian sampling (RMS) trends derived by genomic and traditional evaluations (Tyrisevä et al., 2018a; Tyrisevä et al., 2018b). Genetic selection works by selecting animals with superior Mendelian sampling. The selection is based on phenotypes and progeny records in BLUP, and additionally on genomic information with genomic methods (Lourenco et al., 2020). When some animals are selected for superior Mendelian sampling, the average Mendelian sampling for all the animals is still zero, but for the selected animals is different than zero. Therefore, RMS for genotyped animals is likely to be different than zero with selective genotyping based on performance for both BLUP and ssGBLUP. Additionally, RMS is likely to be zero for both methods when genotyping involves all young animals or is random. However, the magnitude of RMS by ssGBLUP will be bigger because of the higher accuracy of genomic EBV (GEBV). Not only the accuracy is higher, but the average GEBV is usually greater than the average EBV, which translates into superior genetic trends. This study aimed to find the onset of genomic selection by comparing the genetic and Mendelian sampling trends derived by ssGBLUP versus BLUP in pigs, Angus cattle, and broiler chickens.

## Materials and Methods

### Pig data

The pig data consisted of 934,148 records for average daily gain (ADG) and 856,546 for Backfat (BF) collected until 2019, and 1,310,240 animals in pedigree, of which 117,091 were genotyped for 43,910 SNP markers after quality control. This dataset was provided by Genus PIC (Hendersonville, TN). The descriptive statistics of studied traits can be seen in Table 1.

**Table 1.** Descriptive statistics of pig data

| Trait | no. Records | Mean | SD | no. Genotypes | no. Animals in Pedigree |
| --- | --- | --- | --- | --- | --- |
| ADG | 934,148 | 696.86 | 97.45 | 116,943 | 1,310,240 |
| BF | 856,546 | 9.39 | 2.78 | 116,943 | 1,310,240 |

ADG: Average Daily Gain; BF: Backfat; SD: Standard Deviation

### American Angus data

Genotypes, pedigree, and phenotypes for three traits including birth weight (BTW, N=9,003,125), weaning weight (WW, N=9,506,570) and post weaning gain (PWG, N=4,671,702) of Angus beef cattle were provided by the American Angus Association (St. Joseph, MO). The pedigree consisted of 11,573,108 animals, of which 842,199 were genotyped for 39,766 SNP markers. The quality control of genotypes was conducted as in Lourenco et al. (2015b). The descriptive statistics of studied traits in American Angus can be seen in Table 2.

**Table 2.** Descriptive statistics of Angus data

| Trait | no. Records | Mean | SD | no. Genotypes | no. Animals in Pedigree |
| --- | --- | --- | --- | --- | --- |
| BTW (lb) | 9,003,125 | 80.57 | 9.87 | 842,199 | 11,573,108 |
| WW (lb) | 9,506,570 | 593.72 | 99.52 | 842,199 | 11,573,108 |
| PWG (lb) | 4,671,702 | 362.50 | 147.93 | 842,199 | 11,573,108 |
| BTW: Birth Weight; WW: Weaning Weight; PWG: Post Weaning Gain<br>SD: Standard Deviation |  |  |  |  |  |

### Broiler chicken data

The broiler chicken data were provided by Cobb-Vantress Inc. (Siloam Springs, AR). The dataset comprised phenotypes records on a purebred broiler chickens across 32 breeding cycles for three production traits referred as T1, T2 and T3. Each eight breeding cycles comprise one generation. The number of records for T1, T2 and T3 was 1,072,854, 228,992 and 265,891, respectively. The genotype file consisted of 154,318 birds genotyped for 54,713 SNP markers, and the pedigree consisted of 1,252,619 birds. The SNP data underwent quality control process as described in Lourenco et al. (2015a).

### Statistical models

The statistical model for broiler chicken traits was as in Lourenco et al. (2015a), for pig traits was as in Steyn et al. (2020) and for beef traits was as in Garcia et al. (2020). The (co)variance components used in all analyses were provided by Angus Genetics Inc., PIC, and Cobb-Vantress. Both BLUP and ssGBLUP were run in a multiple-trait animal model framework. The pedigree relationship matrix (**A**) was used in BLUP and the realized relationship matrix (**H**) was used in ssGBLUP. The structure of **H^-1^** is explained in Misztal et al. (2009) and Aguilar et al. (2010).

### Genomic analysis and software

Because of the large number of genotyped animals, the algorithm for proven and young (APY) was used to create the inverse of 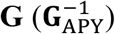 as proposed by Misztal et al. (2014a) and Fragomeni et al. (2015). In APY, the matrix of genomic relationships among genotyped animals is partitioned based on core and noncore animals. The number of core individuals was selected based on the number of eigenvalues explaining 98% of the variance of **G** (Pocrnic et al., 2016) using PREGSF90 (Misztal et al., 2014b). The number of core individuals for broiler chickens, pigs, and beef cattle was estimated as 5030, 11,094, and 13,000, respectively.

Solutions for BLUP and ssGBLUP were obtained by using the preconditioned conjugate gradient algorithm with iteration on data as implemented in the BLUP90IOD2 (Tsuruta et al., 2001). The convergence criterion was set to 10^−12^ for all evaluations.

### Criteria to investigate the starting point of Genomic preselection

#### Genetic trends

The point of divergence in genetic trends obtained by ssGBLUP and BLUP were used as a way to identify the onset of genomic selection. To explain how the difference between predictions from ssGBLUP and BLUP can indicate the start of genomic selection, consider the decomposition of the (genomic) estimated breeding values ((G)EBV) of individual *i* as in Aguilar et al. (2010), VanRaden and Wright (2013), and Lourenco et al. (2015a):

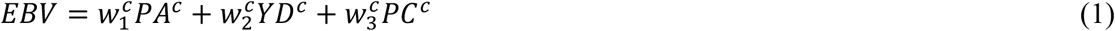

and

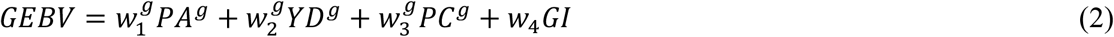

Then, the difference between GEBV and EBV is:

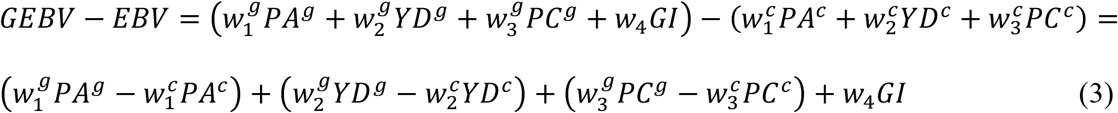

where PA is the parent average, YD is yield deviation (phenotypes adjusted for the fixed effects), PC is the progeny contribution, and GI is the genomic information which is equal to GP-PP, in which GP is the genomic prediction derived using **G** and PP is the pedigree prediction derived using **A**_22_; the superscripts *c* and *g* denote components related to conventional BLUP and ssGBLUP, respectively, and *w*_1_ *to w*_2_ are weights that sum to 1.

When inbreeding is ignored in **A** and both parents are known, then, *w*_1_ = 2/*den, w*_2_ = (*n_rec_/α*)/*den, w*_3_ = 0.5*n_prog_/den*, and 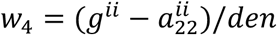, in which *α* is the variance ratio (residual variance over additive genetic variance), *n_prog_* is the progeny size, *n_rec_* is the number of records, *g^ii^*(*a^ii^*) is the diagonal element of 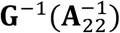 for animal *i, den* is the sum of the numerators of *w*_1_ to *w*_4_.

The components of (G)EBV equations for individual *i* are as following:

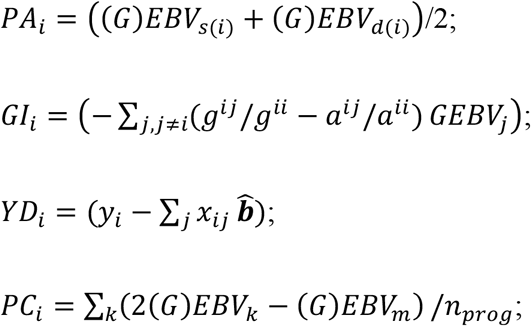

Where (*G*)*EBV_s(i)_* and (*G*)*EBV_d(*i*)_* are (genomic) breeding values of sire and dam of individual *i, y_i_* is the *i*th record of animal *i*, 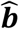 is the solutions for the level of fixed effects related to record *i, x_ij_* is element of a design matrix relating 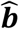 to *y_i_*, and *k* refers to progeny and *m* indicates mate of animal *i*.

The components GP and PP are ignored under BLUP, which results in biased EBV if animals are selected based on genomic information. The bias arises not only from the lack of GP and PP, but from a combination of elements including the fact that PA, PC, and YD are not adjusted based on genomic information. For instance, if parents are non-genotyped, the difference between the predictions from BLUP and ssGBLUP originates from the contributions due to PC and GI of genotyped animals. For young animals without own and progeny records, the difference between EBV and GEBV comes from GI and PA enhanced by genomic information of parents, the latter to a smaller extent. However, as own and progeny records are added to the data, the amount of weight given especially to PC increases, and the weight of GI decreases.

When EBV or GEBV are used for selection of parents, GEBVs have higher accuracy 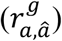. This will generate a difference in amount of genetic gain (Δ*G*) in the next generation. Therefore, it can be shown as 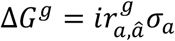 and 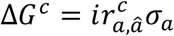, and finally Δ*G^g^* ≥ Δ*G^c^*, in which *i* is the selection intensity and *σ_a_* is the additive genetic standard deviation. Hence, under genomic selection, GEBVs are higher than EBV because greater accuracy of GEBV allows the selection of superior animals based on GP. Subsequently, a divergence in (G)EBV trends indicates the beginning of the genomic selection.

To obtain the genetic trend under traditional BLUP and ssGBLUP, the (G)EBVs were averaged by birth year for genotyped bulls in the beef cattle population and all genotyped individuals in the pig and chicken populations. Only animals with phenotypes were used for deriving the genetic trends. Genetic trends were obtained using a simple linear regression of (G)EBV for each trait on year of birth. For both BLUP and ssGBLUP, the genetic base was set to where more than one thousand genotyped individuals were available per year/generation. This corresponded to breeding cycle 1 in broiler chickens, and birth year 2012 in pigs and, 2007 in beef cattle. The mean GEBV from ssGBLUP was set to the same base as EBV from BLUP.

#### Realized Mendelian Sampling (RMS)

The RMS for the genotyped individual *i* was estimated as:

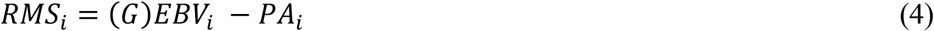

Under some idealized evolutionary process (e.g., random mating, absence of selection, and large population size) all components are expected to be zero for the same generation.

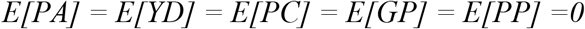

and consequently E[RMS]=0. When all or a random subset of young animals are used as parents of the next generation, the average RMS is close to 0. However, in the population under selection the equalities may not hold; therefore, *E*(*RMS*) ≠ 0.

For simplicity, assume that parents and earlier generations are not genotyped. Let index *s* denotes ungenotyped animals selected for genotyping based on phenotype or BLUP (the first stage of selection), then E[YD]=δ, where 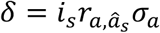, in which *i_s_* is the selection intensity at the first stage of selection, 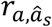 is the accuracy of evaluation based on phenotype or BLUP and *σ_a_* is the additive genetic standard deviation. Assuming young animals with neither progeny nor genotype:

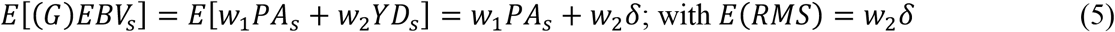

Therefore, if animals are preselected based on phenotype or BLUP, RMS from either BLUP or ssGBLUP is nonzero. Its value depends not only on the selection differential but also on the coefficient *w*_2_, which is a function of variance ratio and the number of records.

Now assume that in the second stage of selection, the animals preselected based on phenotypes or BLUP are genotyped and reevaluated (index *sg*). On average, an animal with superior phenotype may also have a superior genomic prediction, E[GP]=τ, where 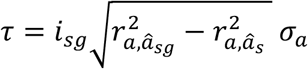, with *i_sg_* selection intensity in the second stage of selection and 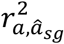 is the reliability of selection based on the genomic reevaluation. Then,

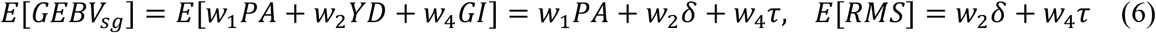

With many genotyped animals, the coefficient *w*_4_ can be close to 1, with accuracy of *GEBV_sg_* greater than the one of *EBV_s_*. Accordingly, RMS will be greater under genomic selection. The selective genotyping based on superior phenotypes (YD) can be replaced by superior progeny difference (PC) indicating that both have a similar effect on EBV, GEBV, and RMS.

The above derivations suggest that the RMS is close to zero when all animals are genotyped or when genotyping is at random. With selective genotyping, RMS is nonzero and is greater with ssGBLUP than with BLUP. Because selective genotyping is the practice in livestock populations, the divergence in RMS trends obtained based on EBV and GEBV can also indicate the start point of the genomic selection. The same animals which were used for obtaining the genetic trends, were engaged in attaining the RMS trends.

## Results

### 1) Pig production traits

Figure 1 shows the genetic trends for ADG and BF in genotyped pigs. The annual changes in average breeding values in genetic standard deviation units from 2012 to 2019 for ADG and BF were 0.27 and 0.04 for ssGBLUP and 0.18 and 0.02 for BLUP, respectively. The trends from ssGBLUP and BLUP diverged after 2013. In the last year of data (2019), the differences between average breeding values from ssGBLUP and BLUP were 0.67 SD for ADG and 0.17 SD for BF.

**Figure 1.**
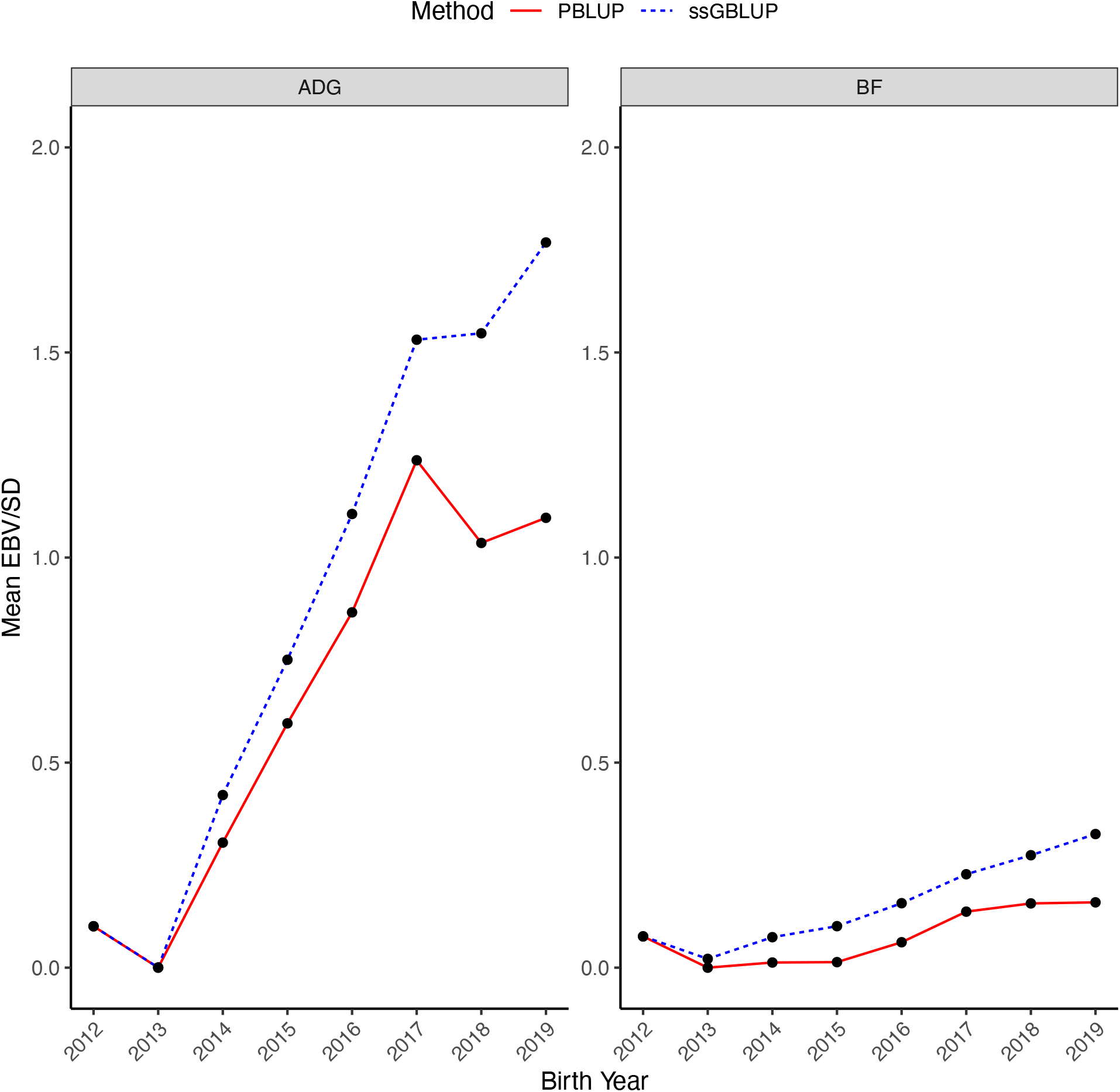
Genetic trends obtained using single-step GBLUP (ssGBLUP) and pedigree BLUP (PBLUP) for average daily gain (ADG) and backfat (BF) in the genotyped pigs by year of birth. Genetic trends are presented in additive genetic standard deviation scale and the genetic base is adjusted to 2012.

The genetic trend for ADG increased over time with a slightly increase in BF observed in recent years. The change in the genetic trend for BF was possibly due to the correlated response with body weight traits, as well as changes in breeding practices and in the selection objective in recent years.

The RMS (Figure 2) for ADG increased from around 0.04 in 2012, reached a peak of 0.10 in 2016, then declined. Relatively large RMS suggests preselection on a correlated trait-before genotyping. Smaller RMS for BF could be due to a correlated response to ADG.

**Figure 2.**
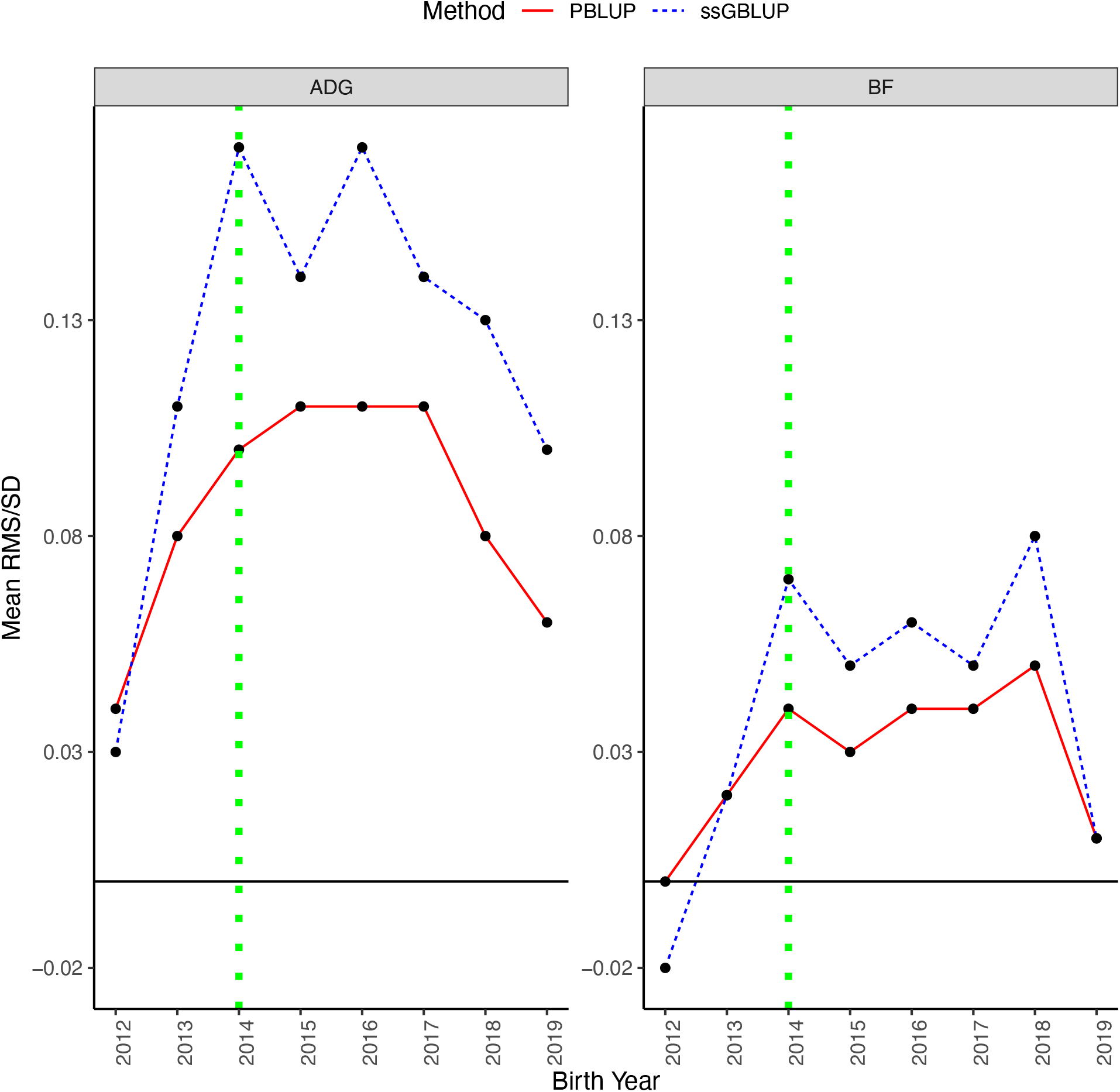
Realized Mendelian sampling (RMS) trends estimated by single-step GBLUP (ssGBLUP) and pedigree BLUP (PBLUP) for average daily gain (ADG) and backfat (BF) in the genotyped pigs. Mendelian sampling trends are presented in additive genetic standard deviation scale. Solid black line represents the zero-base line and dotted green vertical line shows the start date of genomic selection.

### 2) Beef production traits

The genetic trends achieved by BLUP and ssGBLUP for BTW, WW, and PWG in genotyped Angus bulls are shown in Figure 3. The annual changes in (G)EBV for genotyped animals, in genetic standard deviation units, from 2006 to 2018 for BTW, WW, and PWG were −0.01, 0.11, and 0.08 for ssGBLUP and −0.01, 0.09, and 0.09 for BLUP, respectively. In the last year of data (2018), the differences between average breeding values from ssGBLUP and BLUP were 0.01, 0.23, and 0.06 SD for the three traits, respectively.

**Figure 3.**
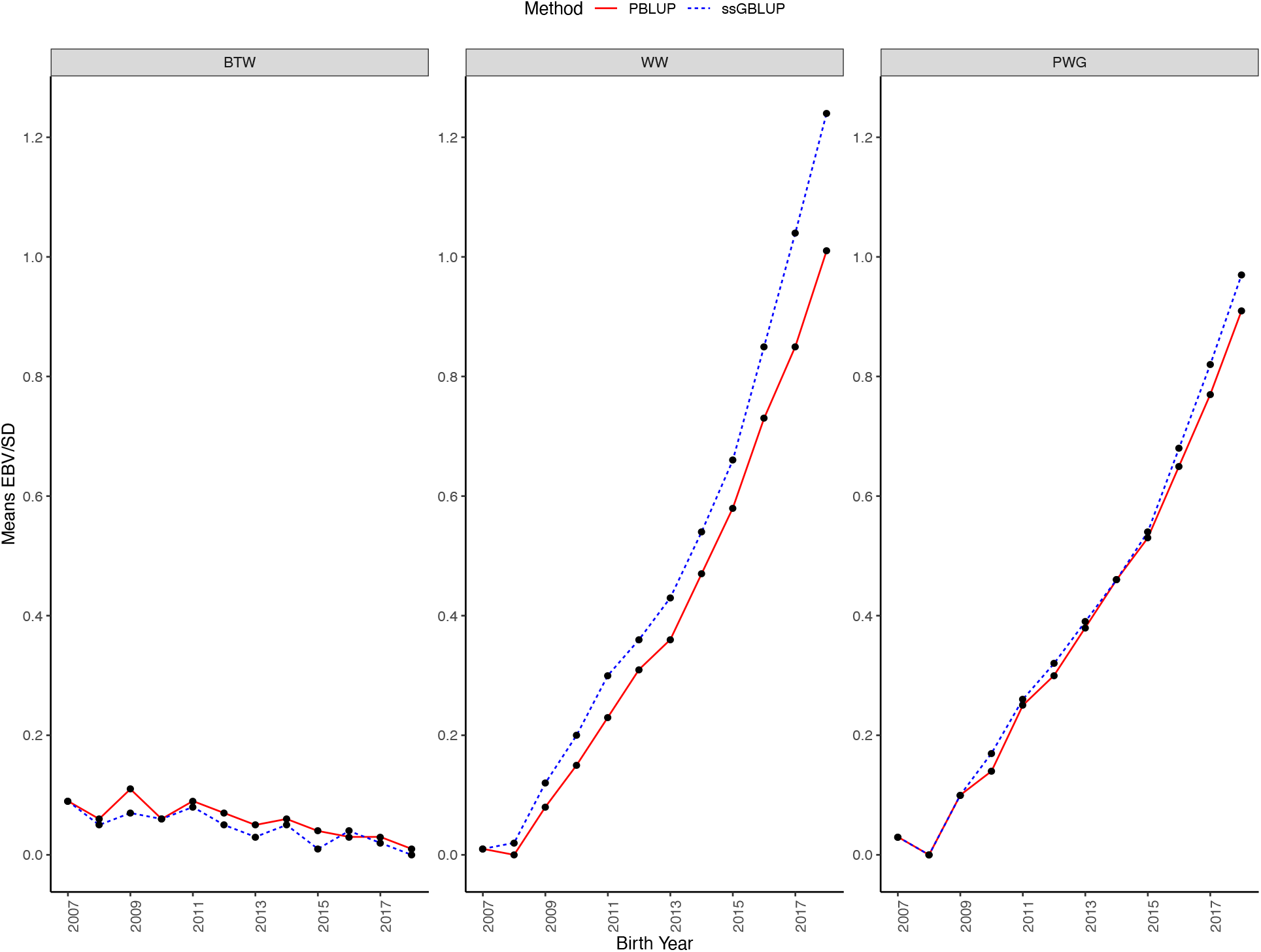
Genetic trends obtained using single-step GBLUP (ssGBLUP) and pedigree BLUP (PBLUP) for birth weight (BTW), weaning weight (WW), and post weaning gain (PWG) in the genotyped Angus bulls by year of birth. Genetic trends are presented in additive genetic standard deviation scale and the genetic base is adjusted to 2007.

For BTW, the difference between the genetic trends for ssGBLUP and BLUP was negligible, but for WW and PWG genetic trends diverged considerably from 2016 afterward. For WW and PWG, the annual genetic gain after 2016 from ssGBLUP was 0.06 and 0.02 SD greater than BLUP, respectively. As it can be seen in Figure 3, there is a genetic improvement for all traits. However, genetic trend of BTW is downward relative to WW and PWG. Low BTW is desirable to avoid calving problems. On the other hand, BTW is positively correlated with WW and PWG; therefore, a stronger pressure is needed to keep BTW low while increasing WW and PWG. Based on the divergence, genomic selection is less important for BTW because this trait has already been recorded at the time of genotyping. Therefore, selection for BTW is based on parent average, phenotype deviation, and genomic prediction. Differently, there was a clear impact of genomic selection for WW from 2009-with an accelerated trend in 2017, and the genomic selection on PWG is slightly visible from 2017.

The RMS (Figure 4) looks very different for the 3 traits. For BTW, the trend is small and negative, at around −0.02, with small changes at the end. It suggests that the heaviest calves were not genotyped; calves are selected for lower BTW to reduce calving difficulty. For WW, RMS is large and increasing over time from 0.12 to 0.29. Such a trend suggests that the primary genotyping is after weaning and based on WW. For PWG, RMS is smaller although rising to 0.17. As the differences between EBV and GEBV were small for PWG, the values of RMS for PWG could be just a correlated response to WW as the genetic correlation between WW and PWG is high.

**Figure 4.**
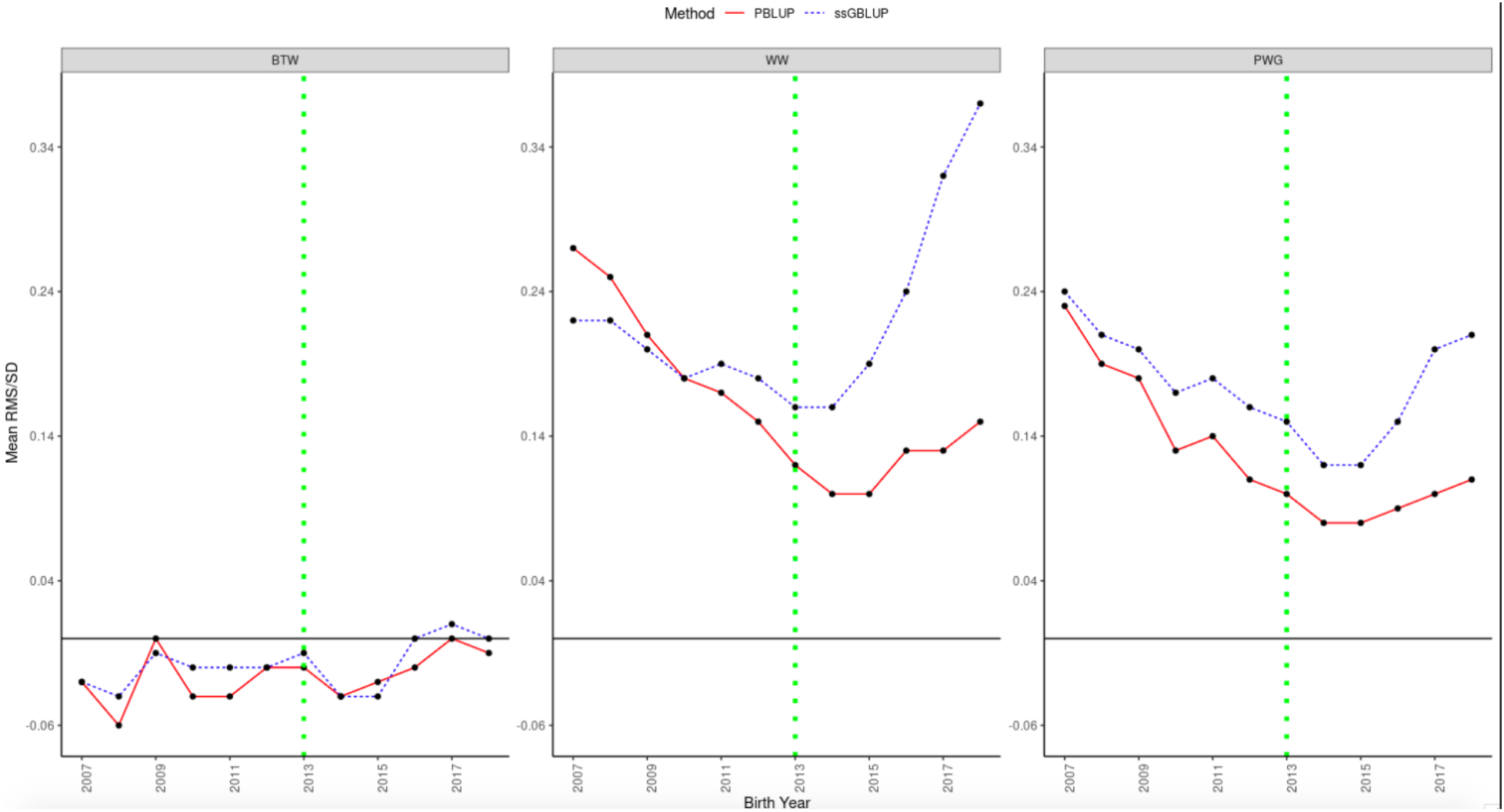
Realized Mendelian sampling (RMS) trends estimated by single-step GBLUP (ssGBLUP) and pedigree BLUP (PBLUP) for birth weight (BTW), weaning weight (WW), and post weaning gain (PWG) in the genotyped Angus bulls. Mendelian sampling trends are presented in additive genetic standard deviation scale. Solid black line represents the zero-base line and dotted green vertical line shows the start date of genomic selection.

### 3) Broiler chicken traits

Trends were favorable for all traits with faster improvement in recent years. Figure 5 shows the difference between genetic trends obtained using ssGBLUP and BLUP in genetic standard deviation units for T1, T2, and T3 in genotyped birds. Divergence for the genetic trends by ssGBLUP and BLUP occurred in breeding cycle 6 for T2 and T3. For T1, some divergence was visible from breeding cycle 2 to 16 in favor of BLUP and then from breeding cycle 20 afterwards in favor of ssGBLUP, although the divergence was reduced later. It seems that for T1 slight divergence in favor of BLUP up to breeding cycle 19 was spurious, and this divergence could represent low genomic merit of animals selected for genotyping.

**Figure 5.**
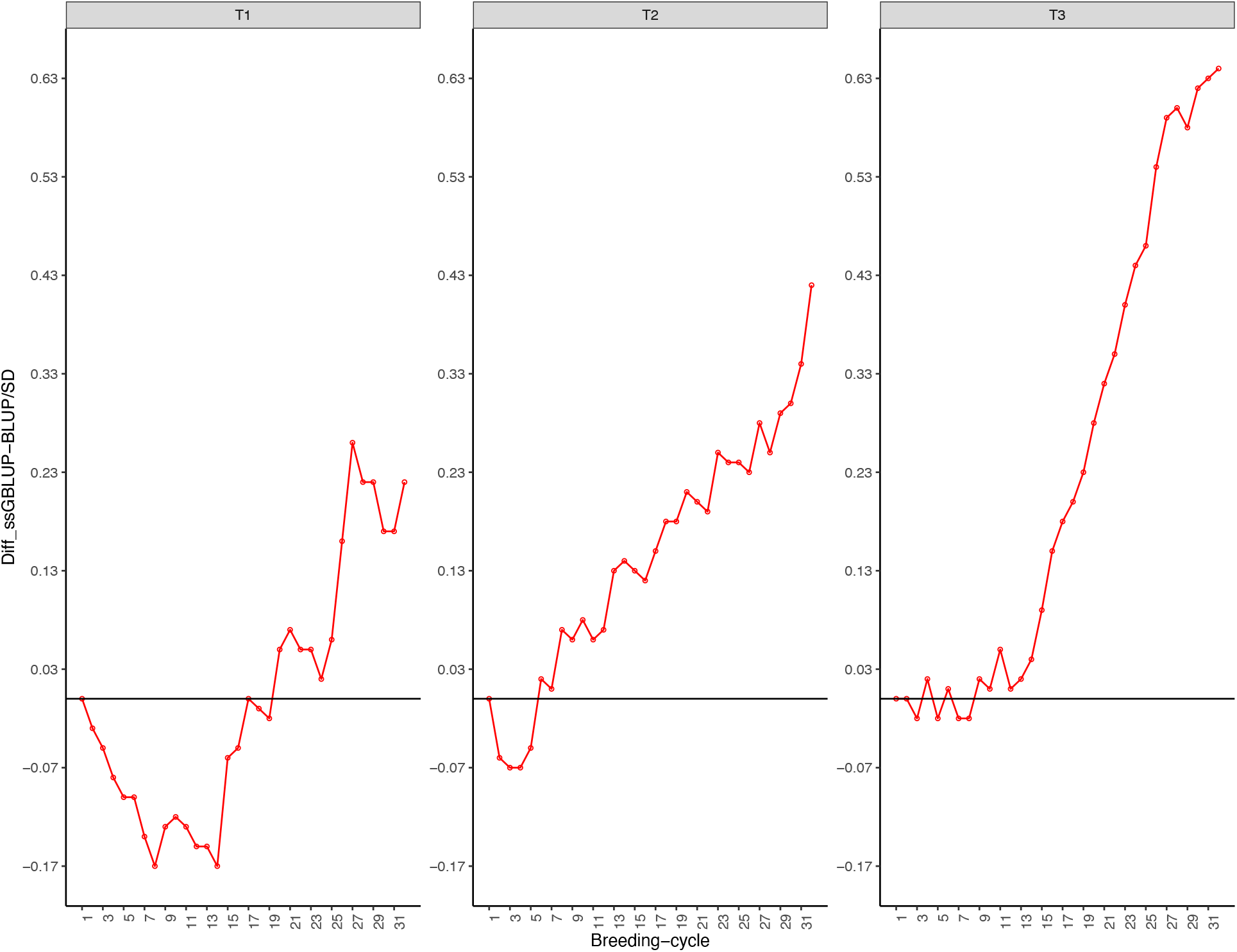
The difference between genetic trends obtained using single-step GBLUP (ssGBLUP) and pedigree BLUP (PBLUP) in genetic standard deviation units for three production traits referred as T1, T2, and T3 in a purebred broiler chicken line across 32 breeding cycles.

The RMS trends (Figure 6) show relatively large values for T1 (up to 0.14) and small values for the other traits (0.04 or less). Animals were selected for T1 by BLUP, then superior animals were genotyped. Therefore, RMS for T1 is high. Small RMS for the other two traits measured later suggests only a correlated response from T1 because all animals measured for these traits were already genotyped.

**Figure 6.**
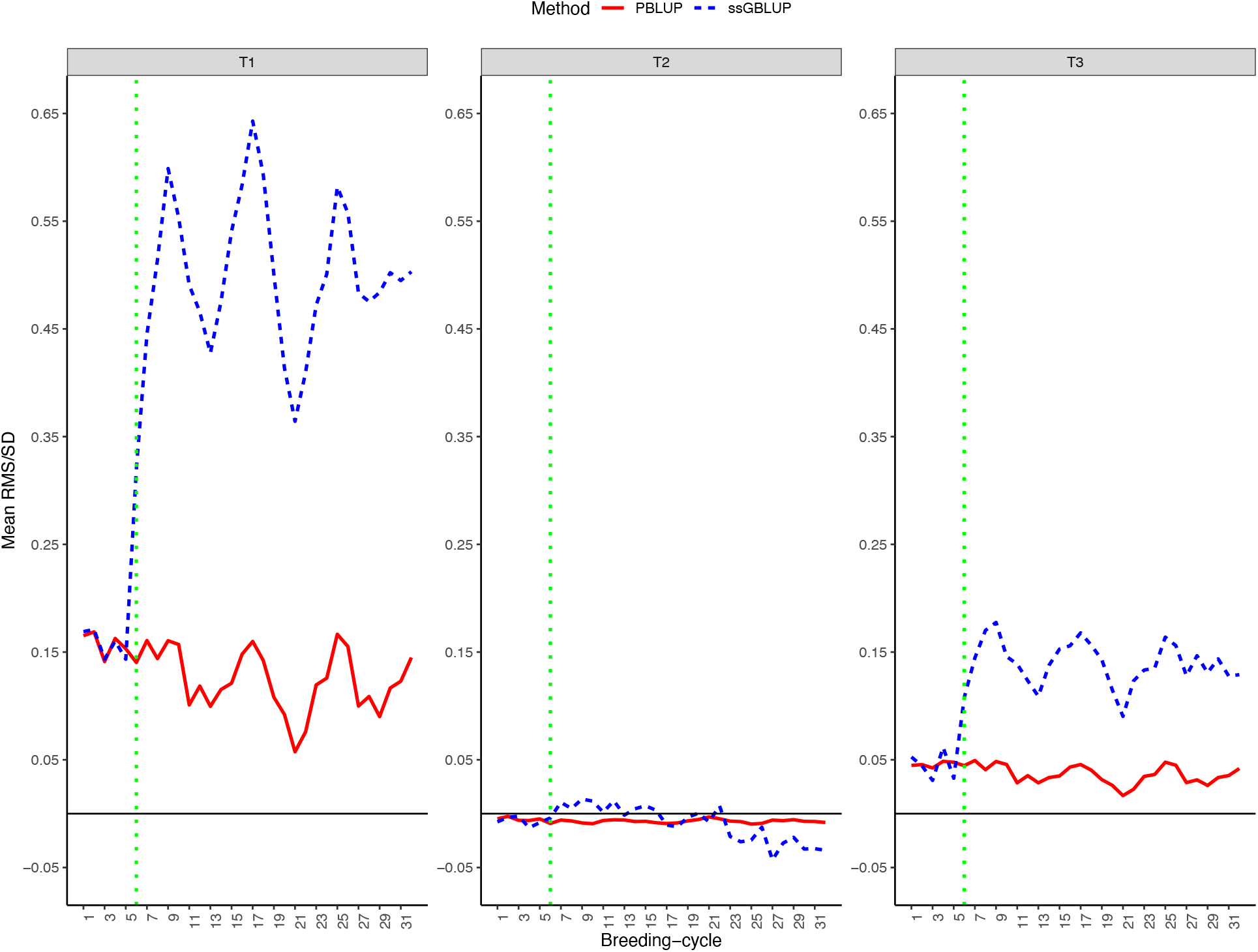
Realized Mendelian sampling (RMS) trends estimated by single-step GBLUP (ssGBLUP) and pedigree BLUP (PBLUP) for three production traits referred as T1, T2, and T3 in a purebred broiler chicken line across 32 breeding cycles. Mendelian sampling trends are presented in additive genetic standard deviation scale. Solid black line represents the zero-base line and dotted green vertical line shows the start date of genomic selection.

## Discussion

### History of adoption of genomic selection

In this study, we used data provided by PIC, American Angus Association and Cobb-Vantress. Although each of them took different approaches when implementing genomic selection and genotyping became available, all changed to ssGBLUP after some time which corresponds to breeding cycle 6 in broiler chickens, year 2014 in pigs and year 2013 in beef cattle.

PIC started using ssGBLUP for genomic evaluations in this population in late 2013, so the first results of genomic selection were visible in 2014. Before that, selection was based on BLUP (William Herring, PIC, Hendersonville, TN, personal communication).

Angus Genetics Inc. incorporated genomic information on 15 markers in 2009 using a correlated trait approach (Kachman, 2008). The panel was updated to 384 markers in 2010 and moved to the 50k SNP chip after that. Finally, ssGBLUP was implemented for Angus cattle evaluations in 2017 (Kelli Retallick, Angus Genetics Inc., St. Joseph, MO, personal communication).

### Genetic trends

We assessed the genetic trends of several traits in broiler chickens, pigs, and beef cattle to investigate the effectiveness of genomic selection. Assuming those differences in genetic basis between BLUP and ssGBLUP are correctly accounted for by the method described in Vitezica et al. (2011), the effectiveness of genomic selection can be evaluated indirectly by measuring the differences between genetic trends from BLUP and ssGBLUP. If the genetic trend by ssGBLUP is accelerating in a favorable direction and the genetic trend by BLUP is decelerating, genomic selection is likely practiced for the particular trait. If the genetic trends by both methods converge to the same point, the selection based on genotypes is not stronger than the selection based on parent average and phenotypes. The genetic trends can also be influenced by the genetic correlations among traits, especially with sequential selection, where a trend for an earlier measured trait influence a trait measured later. Based on the divergence point of genetic trends from BLUP and ssGBLUP in our study, the starting point of genomic selection in Angus cattle is 2013, in pigs is 2014, and in broiler chickens is breeding cycle 6. These starting points agree with the history of implementation of genomic selection in those populations.

If the genetic evaluations are based on ssGBLUP or GBLUP (**H** or **G** matrix), the estimates of genetic trends using BLUP (**A** matrix) are biased provided that a large portion of selected candidates are genotyped. As the correlation between the elements of **G** and **A**_22_ increases, the genetic trends by two methods will converge. However, some factors such as preselection of selection candidates (Jibrila et al., 2020), incomplete pedigree information, and also the existence of young animals without own and progeny records but with genotypic information (Shabalina et al., 2017) makes this difference larger.

The main purpose in investigating genetic trends is to verify whether selection is effective and whether there is an agreement with phenotypic trends. A disagreement suggests changes in the environment, ineffective selection, or biased genetic trends. When there is a disagreement between BLUP and phenotypic trends, but an agreement between the latter and ssGBLUP trends, there is strong evidence for biased BLUP trends. Masuda et al. (2018) showed genetic trends for milk yield traits based on BLUP were biased downwards for US Holstein bulls and cows. Especially for bulls, the bias in EBV was because of failure in accounting for genomic preselection and underestimated PC because daughters were also genotyped, and therefore, preselected before having their phenotypes recorded. In the same study, the authors showed a good agreement between phenotypic and ssGBLUP, meaning the latter can account for preselection and is not biased under genomic selection.

Therefore, when the BLUP trends become biased, it means selection based on genomic information became effective and BLUP EBV–or any measure derived from it, as deregressed proofs–should not be used anymore. It should be noted that not only genomic preselection can cause bias in BLUP evaluations, but also selection on correlated traits (Sorensen and Kennedy, 1984), poorly-defined unknown-parent groups (Misztal et al., 2013), preferential treatments of selection candidates (Dehnavi et al., 2018) and non-random mating (Tsuruta et al., 2020) can generate bias in BLUP.

### Realized Mendelian sampling

The value and trends for RMS illustrate selective genotyping, where the decision to genotype is based on phenotypes or BLUP evaluations. RMS was large for T1 in broiler chickens, for WW in Angus, and for ADG in pigs where genotyping followed phenotyping. That RMS trend indicates that an increasing number of piglets are being genotyped, reducing selective genotyping. As genotyping becomes less expensive while the cost of phenotyping keeps constant, genotyping more young animals becomes economically justified. For broiler chickens, RMS for later traits as T2 and T3 was close to zero, indicating no new preselected genotyping based on these traits.

Although we investigated RMS and genetic trends to identify the starting point of genomic selection, those two approaches are closely related. As genomic selection works by selecting animals with superior Mendelian sampling, there is a sharp increase in breeding values estimated under genomic methods. This increase in breeding values is evident for selected animals and also their progeny (Tyrisevä et al., 2018a), where animals with large number of genotyped progenies are more likely to have greater Mendelian sampling (Masuda et al., 2018). Consequently, because of larger Mendelian sampling, there is an impact in genetic trends when animals are selected based on genomic information, especially if the selection happens before phenotypes are recorded.

## Conclusions

To detect the effective starting point of genomic selection, two possible ways included divergence point of genetic trends and RMS trends obtained by ssGBLUP and BLUP using official datasets from pigs, beef cattle, and broiler chickens were used. The effective starting point of genomic selection in Angus cattle, pigs, and broiler chickens was determined as year 2013, 2014, and breeding cycle 6, respectively. The difference between genetic and RMS trends from ssGBLUP and BLUP is more obvious in a population under more intense selection, as in pigs and broilers compared to beef cattle. In general, the effective starting point of genomic selection can be detected by the divergence between genetic and RMS trends from BLUP and ssGBLUP, although RMS trends are present for traits recorded before genotyping and later used for genotyping decisions. The results and procedures presented here can help to evaluate the efficiency of the implementation of genomic selection in breeding programs.

## List of Abbreviations

BLUP: Best Linear Unbiased Prediction
ssGBLUP: single step Genomic Best Linear Unbiased Prediction
EBV: Estimated Breeding Value(s)
GEBV: Genomic Estimated Breeding Value(s)
SNP: Single Nucleotide Polymorphism
RMS: Realized Mendelian Sampling
APY: Algorithm for Proven and Young
ADG: Average Daily Gain
BF: Backfat
BTW: Birth Weight
WW: Weaning Weight
PWG: Post Weaning Gain
PA: Parent Average
PC: Progeny Contribution
YD: Yield Deviation
GI: Genomic Information

## Acknowledgements

This study was partially supported by Cobb-Vantress (Siloam Springs, AR), Pig Improvement Company (PIC; Hendersonville, TN), Angus Genetics Inc. (St. Joseph, MO), and by Agriculture and Food Research Initiative Competitive Grant no. 2020-67015-31030 from the US Department of Agriculture’s National Institute of Food and Agriculture. We thank Vivan Breen, Rachel Hawken, Ching-Yi Chen, William Herring, Kelli Retallick, and Steve Miller for providing data access. The authors thank Andres Legarra for helpful comments.

## Conflict of interest statement

The authors declare no real or perceived conflicts of interest.

## Notes

### Competing Interest Statement

The authors have declared no competing interest.

